# Stiffening of the Extracellular Matrix is a Sufficient Condition for Airway Hyperreactivity

**DOI:** 10.1101/2020.06.27.175687

**Authors:** Ryan R Jamieson, Suzanne E Stasiak, Samuel R Polio, Jeffrey W Ruberti, Harikrishnan Parameswaran

## Abstract

The current therapeutic approach to asthma focuses exclusively on targeting inflammation and reducing airway smooth muscle force in an effort to prevent the recurrence of symptoms. However, the treatment is not a cure and has little beneficial effect on the progression of asthma. This suggests that there are mechanisms at play that are likely *triggered* by inflammation and eventually become self-sustaining so that even when airway inflammation is brought back under control, these alternative mechanisms continue to drive airway hyperreactivity in asthmatics. In this study, we hypothesized that the stiffening of the airway extracellular matrix is a core pathological change *sufficient* to support excessive bronchoconstriction in asthmatics even when in the absence of inflammation. To test this hypothesis, we increased the stiffness of airway ECM by photo-crosslinking collagen fibers within the airway wall using riboflavin (vitamin B2) and Ultraviolet-A radiation. In our experiments, collagen crosslinking led to a three-fold increase in stiffness of the airway extracellular matrix. This change was sufficient to cause airways to constrict to a greater degree, at a faster rate when exposed to a low dose of contractile agonist. Our results highlight the need for therapeutic approaches that target matrix remodeling to develop a lasting cure for this disease.

## Introduction

Asthma is a debilitating respiratory disease that adversely affects the lives of over 300 million people worldwide (23). Airways in an asthmatic are prone to sudden, exaggerated narrowing in response to a low concentration of inhaled airway smooth muscle (ASM) agonist. A low dose of inhaled agonist that evokes little to no response in a healthy individual, can cause airways to hyper-constrict in an asthmatic. However, the exact mechanisms that trigger a healthy airway to become hypercontractile are not known. There is a consensus in the field that sustained inflammatory signals acting on the ASM play an important role in the development of hypercontractile airways. As such, the treatment of asthma over the past several decades has focused on using a combination of anti-inflammatory drugs, to reduce airway inflammation and bronchodilators which cause relaxation of the ASM. This therapeutic strategy is only aimed at managing the recurrence of symptoms and does not alter the course of progression of the disease(14, 29). Even when inflammation is brought back under control, airways in an asthmatic can still hyper-constrict to a low dose of inhaled contractile agonist (1). This suggests that there are alternative mechanisms at play that are likely *triggered* by inflammation and eventually become self-sustaining. Although airway inflammation can be resolved using anti-inflammatory therapies, these self-sustaining alternative mechanisms may be sufficient to drive airway hyperreactivity (AHR) in asthmatics. In this study, we examined whether an increase in stiffness of the extracellular matrix (ECM) within airways would be *sufficient* to sustain excessive bronchoconstriction in asthmatics even in the absence of inflammation.

From *in-vitro* studies, we know that the ECM plays a critical role in regulating agonist-induced Ca^2+^ oscillations in smooth muscle cells(27). A pathologically stiff matrix can cause healthy ASM cells to falsely perceive a higher dose of contractile agonist than they actually received. Similarly, mechanobiological interactions between the ASM and a stiff environment have been shown to lead to ASM hypercontractility (3, 24), ASM hyperplasia (26) and remodeling that continues unabated by bronchodilators or glucocorticoids (8): all of which are characteristic features of asthma. Despite the wealth of data at the cellular level, the potential role of ECM stiffening in driving the development of AHR in asthma has never been tested experimentally at the level of an airway. Among the multitude of proteins, proteoglycans, and glycoproteins that make up the airway ECM, collagen is the most critical determinant of ECM stiffness and failure strain in the lung (17). In this study, we increased the stiffness of airway ECM by crosslinking collagen fibers within the airway wall using riboflavin (vitamin B2) and Ultraviolet-A radiation (UV-A). This collagen crosslinking procedure is a well-established method for increasing ECM stiffness in multiple tissue types (31, 35, 36), and is routinely used in a clinical setting to increase the stiffness of collagen within the human cornea (35).

We found that a healthy airway can be made hyperreactive simply by increasing the stiffness of the ECM in the airway wall. Our data demonstrate that any remodeling that increases the stiffness of the airway ECM is *sufficient* to drive AHR, even in the absence of inflammatory triggers. Our results highlight the need for therapeutic approaches that target ECM remodeling to develop a lasting cure for this disease.

## Materials and Methods

### Tissue preparation

Bovine lungs were obtained from a local slaughterhouse (Research 87, Boylston, MA) immediately after death, placed on ice, and delivered to our lab within a few hours. A bronchus of the right lung (generations 10–15) was dissected free of parenchymal tissue, and bronchial rings approximately 1-3 mm in diameter were cut from the bronchus for imaging. Tissue specimens were kept in a heated water bath at 37°C in warmed Krebs solution (in mM: 118 NaCl; 4.7 KCl; 1.2 MgSO_4_·7H_2_O; 25 NaHCO_3_; 1.2 KH_2_PO_4_; 11 Glucose; and 2.5 CaCl2). Tissue viability was then confirmed by subjecting the bronchial rings to electric field stimulation (EFS)(7, 15). The airway tissue samples used in this study came from 3 separate bovine lungs. This study was carried out in accordance with the guidelines and regulations approved by the Institutional Biosafety Committee at Northeastern University.

### Measurement of airway reactivity

Viable bronchial rings were subjected to a dose of 10^−5^M acetylcholine (ACh) and allowed to freely constrict. The constricting bronchial rings were imaged at a rate of 10 frames per second using a tabletop microscope (Dino-Lite Edge AM7115 Series, AnMo Electronics Corp., Taiwan). Two separate videos were captured. The first video corresponded to a 30-second recording of the airway at baseline. The airway was then exposed to ACh and the constricting airway was imaged for 5 minutes, which is the typical time needed for an airway to fully constrict in response to ACh (15). The resulting image sequences were analyzed using a custom MATLAB image processing algorithm which outlines the inner lumen of the airway in each frame. The algorithm returns the inner luminal area of the constricting airway as a function of time. The time t=0 corresponds to the time at which the airway is first exposed to ACh. The inner luminal area of the airway segment at baseline (*A*_*baseline*_) was calculated by averaging area measurements taken during the last 5 seconds of the baseline video recording. Area measurements collected during the last 5 seconds of the constriction video recording were averaged to calculate the final constricted inner luminal area (*A*_*constricted*_). From these two measurements, we calculate the reactivity of an airway, ***ξ*** as the fractional change in inner lumenal area of the airway

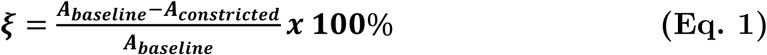

### Increasing the stiffness of the airway ECM

Here, we crosslink collagen fibers using riboflavin (vitamin B2) and UV-A to increase the stiffness of the airway ECM. The wavelength of UV-A light used for these experiments was 365 nm. Collagen crosslinking with riboflavin and UV-A treatment is a well-established technique used to increase ECM stiffness in multiple tissue types (16, 31, 35, 36), and is commonly used in human corneal surgery to increase the stiffness of collagen in the corneal ECM. Once the pre-treatment airway reactivity was recorded, airways were washed 5 times to remove any remaining ACh and allowed to return to their baseline (unconstricted) state. The airways were then immersed in a 1% (w/v) solution of riboflavin (R9504, Sigma-Aldrich) diluted in warm Krebs for 5 minutes. Following riboflavin treatment, the airways were moved to fresh, warmed Krebs solution followed by 15 minutes of UV-A exposure (Blak-Ray Lamp, Model XX-15BLB, 115V, 60 Hz, 0.68 Amps). The intensity of the UV-A light used for experiments was measured at 2.69 mW/cm^2^ using a UV intensity meter (OAI Instruments, Milpitas, CA, Model 0308-0000-01). After 15 minutes of UV-A exposure, the tissue is again washed 5 times in fresh, warmed Krebs solution and tested for viability using EFS. None of the airway samples we used in our experiments failed this viability test. Using the same procedure described in the previous subsection, we then measured airway reactivity (**Eq. 1**). The entire protocol used for collagen crosslinking is summarized in **Fig. 1**.

**Figure 1:**
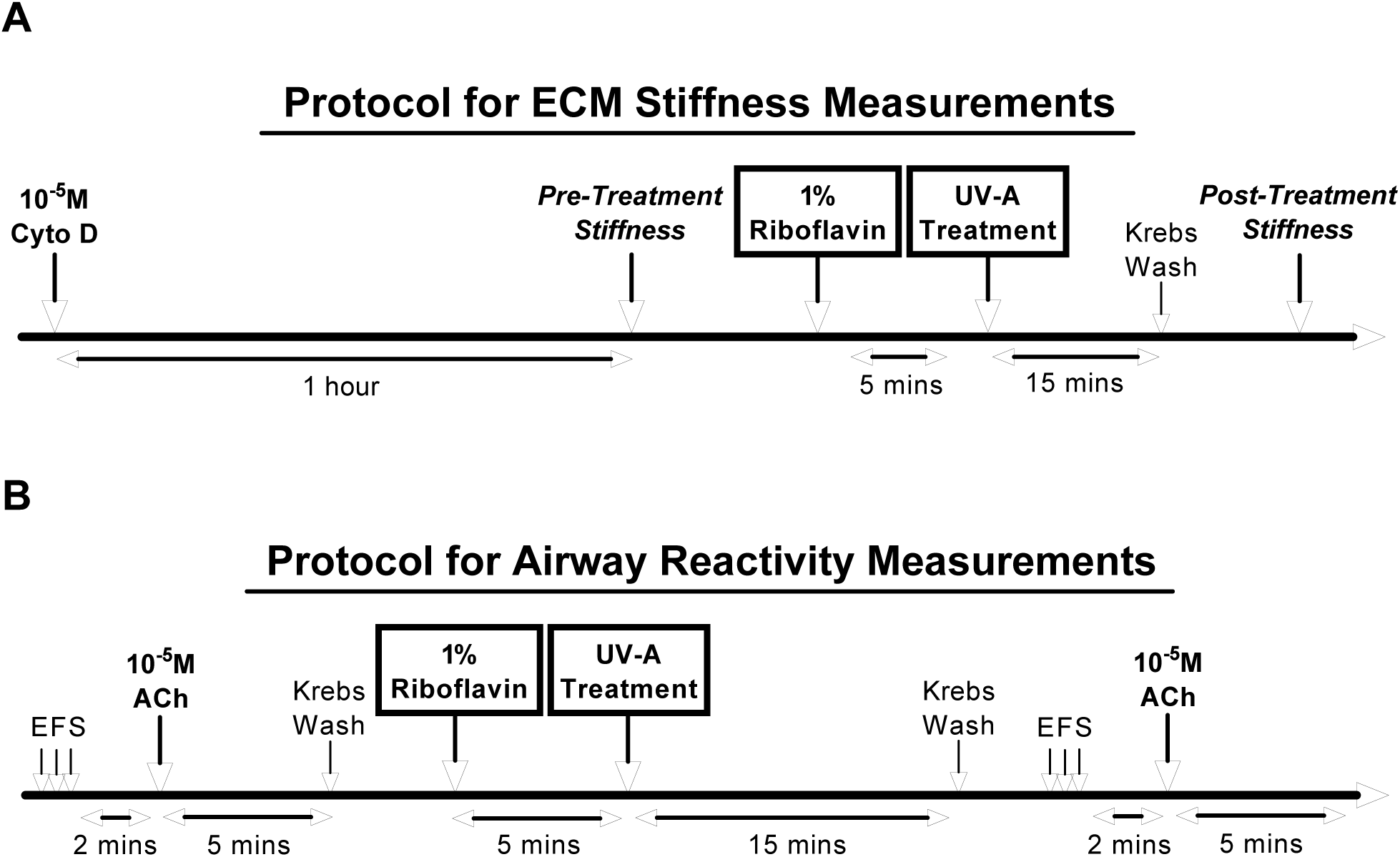
Schematic showing the experimental protocol for **(A)** measuring the ECM stiffness of airways before and after crosslinking the collagen and **(B)** Measuring the reactivity of airways to a fixed dose of contractile agonist (10^−5^M ACh).

### Measurement of ECM stiffness in airways

To measure ECM stiffness, the airways were first treated with 10^−5^M cytochalasin D for 1 hour. Cytochalasin D is a specific inhibitor of actin polymerization in cells, which acts on the positive ends of actin filaments to prevent both the association and dissociation of actin monomers (9). One hour of cytochalasin D treatment disrupts the actin cytoskeleton and significantly decreases the contribution to total tissue stiffness (2). The stiffness of airways measured following cytochalasin D treatment represents the upper bound of airway ECM stiffness. A detailed description of why we chose to measure ECM stiffness in this manner over alternative methods can be found in the Discussion section.

After 1 hour of cytochalasin D exposure, airway segments were mounted in a uniaxial tissue stretcher setup (model 300C, Aurora Scientific, Ontario, Canada) to measure the Young’s modulus of the tissue. A detailed description of the protocol and our experimental setup can be found in Polio et al. (24). Briefly, after preconditioning each airway segment to eliminate prior stretch history, each airway segment was stretched in the radial direction to a maximal strain of 80-100%, then back to 0% using a triangular displacement waveform lasting 60 seconds. The strain was calculated using the length imposed by the lever arm, (***d*)**, and the undeformed inner diameter of each airway segment, (***d***_0_), using **Eq. 2**.

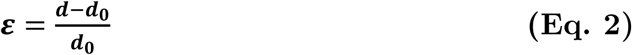

Following force measurements, the longitudinal length (***l***) and wall thickness (***w***) of each airway segment were measured. The force (***F*)**, generated by the stretching of the airways after 1 hour of cytochalasin D treatment was used to calculate airway wall stress, (***σ***), using **Eq. 3**.

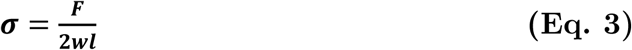

Once the stress-strain curve was generated for each airway, the Young’s Modulus (***E***) was calculated at 60% radial strain by calculating the slope of the stress-strain curve using **Eq. 4**.

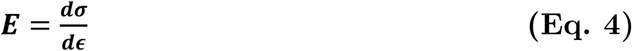

Pre-treatment measurements represent the ECM stiffness of airways treated with cytochalasin D for 1 hour before riboflavin and UV-A treatment, and post-treatment measurements represent the same measurements collected after riboflavin and UV-A treatment.

### Statistical testing

Sigmastat (Systat Software, San Jose, CA) was used to perform statistical testing of experimental data. The specific tests used to determine statistical significance, the number of samples, and the p-value are described along with the corresponding results. Data in the results section is presented as mean ± standard deviation. A p-value of 0.05 was used as a threshold for a statistically significant difference between data sets.

## Results

### Crosslinking of collagen with riboflavin and UV-A increases the stiffness of the airway ECM

A typical stress-strain curve of an airway exposed to cytochalasin D before (pre-treatment) and after (post-treatment) riboflavin and UV-A treatment is shown in **Fig. 2A**. The contribution of cells to the measured stress in **Fig. 2A** is negligible since cells mostly derive their stiffness from cytoskeletal prestress (32). The stress that develops in the airway tissue is predominantly due to the extracellular components in the airway wall (primarily collagen).

**Figure 2.**
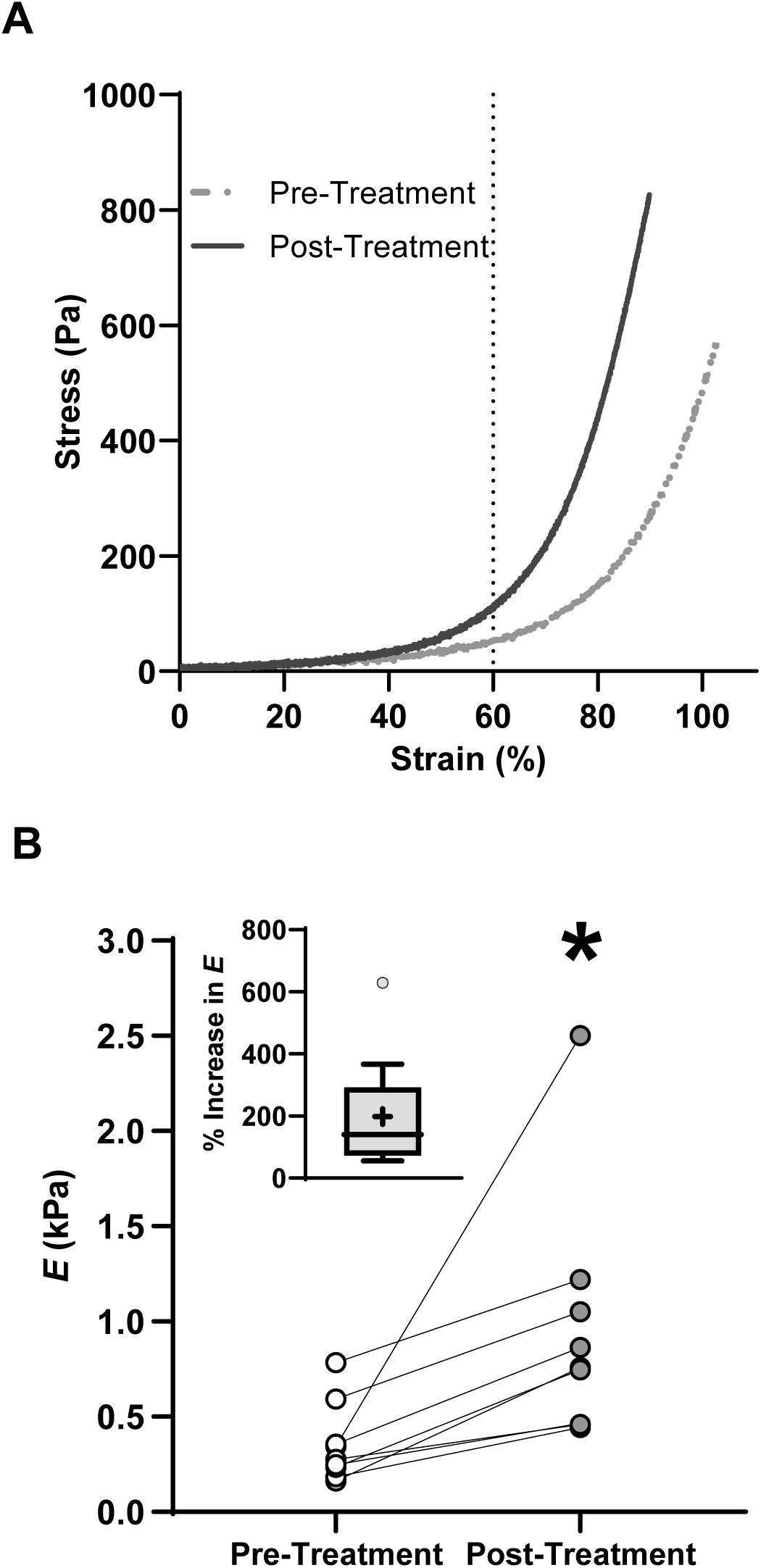
**(A)** Stress-strain curves of an airway ring that has been exposed to 10^−5^M Cytochalasin D for one hour before (dotted grey line) and after (solid black line) treatment with Riboflavin +UVA to crosslink collagen. The Young’s modulus, E was calculated as the slope of the stress-strain curve at 60% radial strain. Exposure to cytochalasin reduces the contribution of the cells to the total tissue stiffness to negligible levels. The value of E measured here represents ECM stiffness. **(B)** Following collagen crosslinking, E increased for all N=9 samples we tested. The effect is statistically significant (N =9, One-way rm-ANOVA, P=0.004). The % increase in E is shown in the inset.

The Young’s Modulus, (***E***), of the airway ECM, increased following riboflavin and UV-A treatment (**Fig. 2B**). Due to a lack of normality in the pre-treatment and post-treatment stiffness values, a repeated measure one-way ANOVA test on ranks was performed. The results of the test demonstrated a significant increase in (*E*) from **353**.**6±205**.**9 Pa** (pre-treatment) to **945**.**5±643 Pa** (post-treatment, N=9, P=0.004). Pre-treatment median of E was 275.3Pa and the post-treatment median was 758.8 Pa). The increase in stiffness due to riboflavin and UV-A treatment was highly reproducible, and all the treated airways demonstrated an increase in ECM stiffness. Across all the samples tested, the mean % increase in ECM stiffness following riboflavin and UV-A treatment was 198.1% (**Fig. 2B inset**).

### Collagen crosslinking leads to a higher degree of airway constriction in response to the same dose of contractile agonist

To measure agonist-induced airway constriction, we first applied a 10-second EFS pulse train to test the viability of the airway tissue (**Fig 1**). EFS pulses cause nerve endings in freshly dissected airways to release ACh, causing the airway to constrict (5, 33). All of our airway samples constricted in response to EFS pulses both before and after riboflavin and UV-A treatment. The mean reactivity of airways in response to EFS before treatment was **75**.**4±15**.**2%**, and **86**.**6±14**.**0%** after treatment with riboflavin and UV-A, indicating that these airways were viable throughout the course of our experiments.

To measure the change in reactivity to a *fixed, low* dose of contractile agonist, we exposed our airway samples to 10^−5^M ACh. We found that exposure to this concentration of ACh caused a low to moderate constriction in our untreated airway samples (***ξ* =36**.**2±37**.**4%**, median ***ξ*=23**.**7%**, N=9). The distribution of ***ξ*** in our untreated samples **(Fig. 3C)** was skewed towards low numbers, so for the samples we tested, 10^−5^M ACh could be considered as a “low” dose of agonist. The degree to which the airways constricted to the same low dose of agonist (10^−5^M ACh) changed significantly after collagen was crosslinked in the same airway samples. The time course of change in the inner luminal area of an airway after adding 10^−5^M ACh, before and after collagen crosslinking is shown in **Fig. 3A**. The baseline and constricted inner luminal area of each airway sample (N=9) before, and after collagen crosslinking is shown in **Fig. 3B**. After collagen crosslinking, airways exposed to the same low dose of ACh (10^−5^M) exhibited a substantial increase in reactivity from ***ξ* =36**.**2±37**.**4%**, N=9, pre-treatment to ***ξ*= 74**.**5±28**.**7%**, N=9 (post-treatment) (**Fig. 3C**). To test for statistical significance, a one-way repeated measure ANOVA test was performed with riboflavin and UV-A treatment as the independent factor, and airway reactivity ***ξ*** measured before and after treatment as the dependent variable. We found that collagen crosslinking led to a statistically significant increase in airway reactivity ***ξ*** for the same dose (10^−5^M) of contractile agonist (ACh). (P ≤0.001, One-way RM ANOVA, N=9).

**Figure 3.**
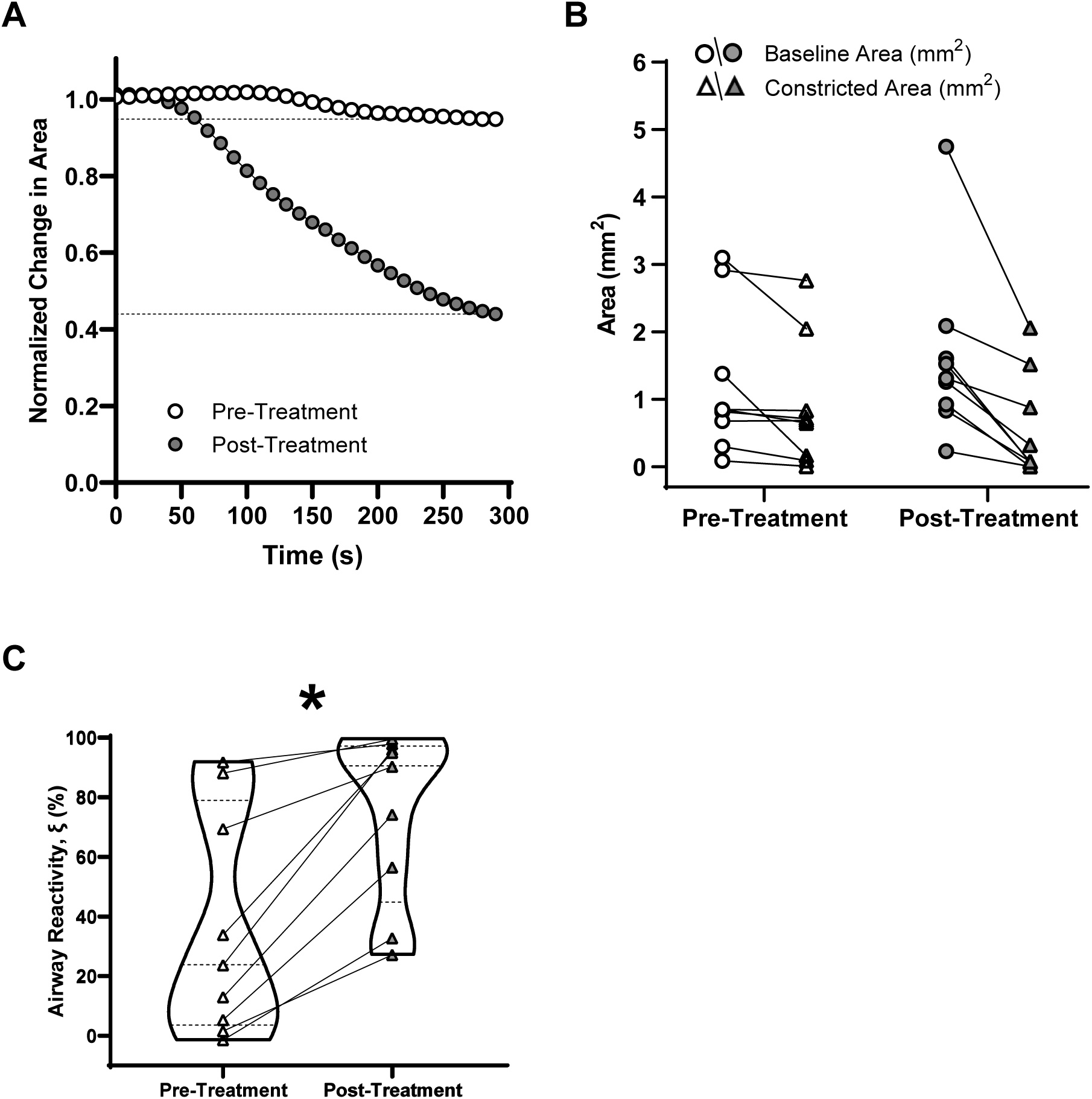
**(A)** Time course of change in the inner luminal area of an airway after adding 10^−5^M ACh, before and after collagen crosslinking. **(B)** The baseline and constricted inner luminal area of each airway sample (N=9) before, and after collagen crosslinking. **(C)** Truncated violin plots of reactivity, ξ. After collagen crosslinking, airways exposed to the same low dose of ACh (10^−5^M) exhibited a substantial increase in reactivity from ξ =36.2±37.4%, N=9, pre-treatment to ξ= 74.5±28.7%, N=9 (post-treatment). The effect is statistically significant (P ≤0.001, One-way RM ANOVA, N=9).

The variable ***ξ*** represents the fractional change in the inner luminal area of an airway in response to a contractile agonist. The use of fractional change in luminal area is the most appropriate means of measuring the response of airways to agonist and is widely used in the literature (4, 30). However, the mathematical formulation of fractional measures makes them prone to bias due to differences in the baseline measure. In our case, ***ξ*** is sensitive to change in *A*_*baseline*_ before and after treatment with riboflavin and UV-A. A lower value of *A*_*baseline*_ post-treatment can lead to higher percent changes in the luminal area and confound our interpretation of the degree of airway reactivity, ***ξ***. To test for the confounding effect of variations in *A*_*baseline*_, we tested for statistical differences in the baseline (pre-ACh) inner luminal area of our airway samples before and after collagen crosslinking and found no difference in *A*_*baseline*_ before and after collagen crosslinking (P= 0.261, One-way RM ANOVA N=9). In the supplementary information, we include two movies of the same airway narrowing in response to ACh before and after treatment with riboflavin and UV-A.

### Collagen crosslinking increases the rate of airway constriction

To test how riboflavin and UV-A treatment affected the rate of airway constriction, the slope of a line that best fits the decrease in the inner luminal area of each airway was measured after exposure to 10^−5^M ACh. Inner luminal area measurements were averaged over 10-second intervals, and constriction rates were calculated for airways before and after treatment. An example time course for a constricting airway after treatment, along with the best-fit line used to calculate the rate of constriction for the airway, is shown in **Fig. 4A**. The pre-treatment and post-treatment rates of constriction for each airway are displayed in **Fig. 4B**. Every sample we tested demonstrated a faster rate of constriction in response to ACh after treatment with riboflavin and UV-A (**1**.**28 ± 1**.**6×10^−3^ mm^2^/s**, pre-treatment vs **5.19±3**.**1×10^−3^ mm^2^/s**, post-treatment). The difference in constriction rates due to collagen crosslinking was statistically significant (P = 0.006, One-way RM ANOVA, N =9).

**Figure 4.**
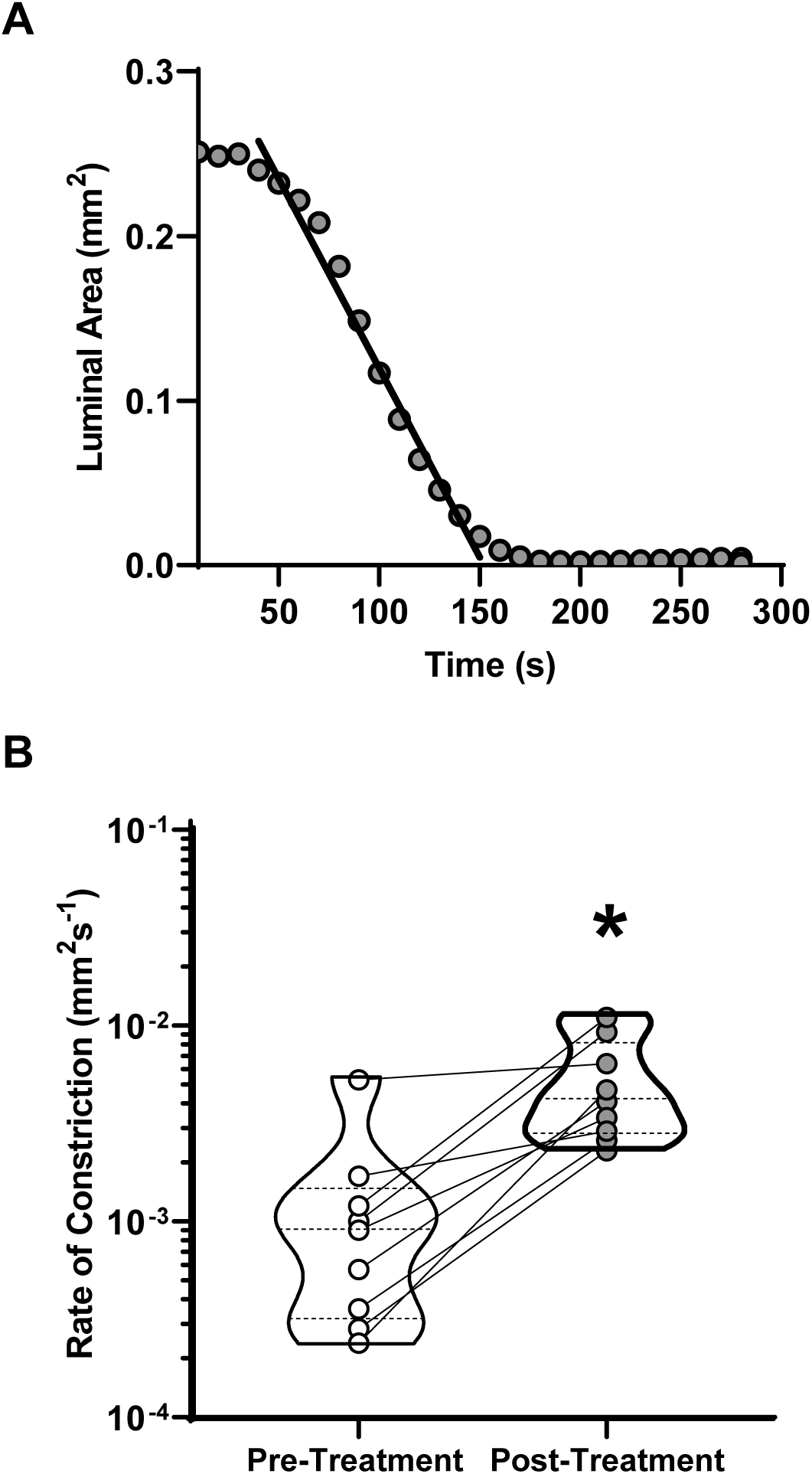
**(A)** Inner luminal area measurements were averaged over 10-second intervals, and constriction rates were calculated for airways before and after treatment. Figure shows an example time course for a constricting airway after treatment, along with the best-fit line used to calculate the rate of constriction for the airway **(B)** The pre-treatment and post-treatment rates of constriction for each airway are shown. The difference in constriction rates due to collagen crosslinking was statistically significant (P = 0.006, One-way RM ANOVA, N=9).

## Discussion

Disruption of collagen homeostasis and excessive deposition of fibrillar collagen are characteristic structural changes found in airways of asthmatics (9, 22). While these pathological alterations in the ECM are likely triggered by inflammation, these changes in composition and rigidity of the extracellular component of the airway wall persist even when airway inflammation is brought under control. In this study, we hypothesized that increasing the stiffness of ECM within the airway wall is a pathological change sufficient to support excessive bronchoconstriction in asthmatics, even in the absence of inflammatory signals. To test this hypothesis, we employed a collagen crosslinking technique using riboflavin (vitamin B2) and UV-A light, which has been shown to increase the stiffness of the ECM within multiple tissue types (31, 32, 35). In our experiments, we were able to generate a three-fold increase in the ECM stiffness of bovine airways (**Fig. 2B**). The goal here was not to recreate the pathophysiological process that occurs in asthma, but to perturb the mechanical properties of the ECM without introducing any inflammatory signals to examine its impact on agonist-induced airway constriction. When we measured the degree of airway reactivity after ECM stiffening, we found that the airways constricted to a greater degree (**Fig. 3C**) and at a faster rate (**Fig. 4B**) in response to the *same dose* of contractile agonist. These results provide direct evidence that structural changes in the wall of healthy airways which result in a pathologically stiffer ECM are sufficient to sustain airway hyperreactivity, even in the absence of inflammation.

The role of collagen remodeling in asthma has been overlooked thus far because of the significant increase in ASM mass seen in severe asthmatics (18). However, in severe asthma, the volume of ECM within the airways is significantly increased compared to healthy and mild asthmatics (18). Further, the fractional area occupied by the ECM within the ASM layer decreases significantly in severe asthma. The reduced volume fraction has been attributed to pathological collagen remodeling, which results in collagen being packed into dense bundles of aligned fibers which occupy less volume (28). Mechanically, such dense, aligned collagen fiber networks invariably possess a high Young’s modulus (20, 21). This reasoning leads us to conclude that the ASM cells in severe asthmatics exist in a very stiff extracellular environment. From *in-vitro* studies, we know that ASM cells become hypercontractile (26), and generate more force for a given dose of agonist (3) when cultured in stiff environments which mimic remodeled ECM. Despite clear evidence at the cellular level, the role of collagen stiffening in driving the development of AHR in asthma has never been experimentally tested at the airway level.

In this study, we measured ECM stiffness by measuring the Young’s modulus of the airway tissue after 1 hour of exposure to cytochalasin D. Cytochalasin is a cell-permeable toxin that acts as potent inhibitor of actin polymerization. By binding to the barbed (+) end of the actin filaments, cytochalasin prevents addition of actin monomers (11). The half-life of actin turnover in cells is approximately 30 seconds (34). Therefore, treatment with cytochalasin D over the course of an hour is enough to disrupt the actin cytoskeleton within ASM cells (2). Since cell stiffness is primarily determined by cytoskeletal prestress, the contribution of cells to the total tissue stiffness in our measurements is negligible (32). So, the measurements shown in **Fig. 2B** represent the upper bound of ECM stiffness in airways. An alternative method for measuring ECM stiffness would have been to decellularize the tissue. While this approach completely removes all the cells from the tissue, there is no decellularization protocol that can perfectly preserve the ECM in tissue. In contrast, the activity of cytochalasin D is specific to actin in the cells. Treatment with cytochalasin is not known to affect ECM components. Regardless of the technique used to abrogate the contribution of cells to the total stiffness of the airway, the stiffness of the tissue should not be seen as a linear function of the cell and ECM stiffness. *In-situ*, the cells attach and pull on nonlinearly elastic components of the ECM, such as collagen, increasing the stiffness of the ECM. At the moment, it is extremely challenging to account for this interaction in estimating ECM stiffness using established experimental techniques.

The stiffness of airway ECM in healthy human lungs is size-dependent. Small airways with an inner diameter ≤ 3mm, which are known to collapse in asthma, possess very soft ECM with a Young’s modulus on the order of 1 kPa (24). If there is a triggering event, perhaps due to a brief period of airway inflammation, collagen is excessively deposited and remodeled within the ASM layer of an airway (13, 25). We recently demonstrated that an increase in matrix stiffness can cause ASM cells to lose direct cell-cell connections and form focal adhesions with collagen fibers in the underlying ECM. This ECM stiffness-induced switch in connectivity causes cells to generate higher forces, which are then transmitted through the collagen fibers (24). This change in the pathway of force transmission through collagen has two important and unexplored consequences for asthma progression. First, collagenase enzymes become ineffective at cleaving collagen fibers that carry tension (6, 10, 12), leading to a further increase in collagen in the ASM layer. Second, due to the strain stiffening properties of collagen, increased force transmission through fibers further increases collagen stiffness (19). Together, these two mechanisms can create a self-sustaining feedback loop that can promote pathological collagen remodeling and AHR development *in the absence of sustained airway inflammation*. This process is entirely driven by mechanical factors and can persist even when inflammation is brought under control. Thus, collagen remodeling in the airway can continue unabated by anti-inflammatory therapy (8).

Our results show that alterations in the mechanical properties of the ECM alone can sustain AHR even in the absence of inflammatory signals, causing the disease to progress unaffected by current therapies. These results highlight the need to therapeutically target ECM remodeling to provide a lasting cure for this debilitating disease.

## Notes

### Competing Interest Statement

The authors have declared no competing interest.

